# Assessing real-time Zika risk in the United States

**DOI:** 10.1101/056648

**Authors:** Lauren A Castro, Spencer J Fox, Xi Chen, Kai Liu, Steve Bellan, Nedialko B Dimitrov, Alison P Galvani, Lauren Ancel Meyers

**Affiliations:** Department of Integrative Biology, The University of Texas at Austin, Austin, TX, USA; Graduate Program in Operations Research Industrial Engineering, The University of Texas at Austin, Austin, TX, USA; Institute for Cellular and Molecular Biology, The University of Texas at Austin, Austin, TX, USA; Center for Computational Biology and Bioinformatics, The University of Texas at Austin, Austin, TX, USA; Center for Infectious Disease Modeling and Analysis, Yale School of Public Health, New Haven, CT, USA; Department of Ecology and Evolution, Yale University, New Haven, CT, USA; The Santa Fe Institute, Santa Fe, NM, USA

**Keywords:** Zika, ZIKV, importation risk, autochthonous transmission risk

## Abstract

**Background:** Confirmed local transmission of Zika Virus (ZIKV) in Texas and Florida have heightened the need for early and accurate indicators of self-sustaining transmission in high risk areas across the southern United States. Given ZIKV’s low reporting rates and the geographic variability in suitable conditions, a cluster of reported cases may reflect diverse scenarios, ranging from independent introductions to a self-sustaining local epidemic.

**Methods:** We present a quantitative framework for real-time ZIKV risk assessment that captures uncertainty in case reporting, importations, and vector-human transmission dynamics.

**Results:** We assessed county-level risk throughout Texas, as of summer 2016, and found that importation risk was concentrated in large metropolitan regions, while sustained ZIKV transmission risk is concentrated in the southeastern counties including the Houston metropolitan region and the Texas-Mexico border (where the sole autochthonous cases have occurred in 2016). We found that counties most likely to detect cases are not necessarily the most likely to experience epidemics, and used our framework to identify triggers to signal the start of an epidemic based on a policymakers propensity for risk.

**Conclusions:** This framework can inform the strategic timing and spatial allocation of public health resources to combat ZIKV throughout the US, and highlights the need to develop methods to obtain reliable estimates of key epidemiological parameters.

## Background

In February 2016, the World Health Organization (WHO) declared Zika virus (ZIKV) a Public Health Emergency of International Concern [1]. Though the Public Health Emergency has been lifted, ZIKV still poses a great threat for reemergence in susceptible regions in seasons to come [2]. In the US, the 268 reported mosquito-borne autochthonous (local) ZIKV cases occurred in Southern Florida and Texas, with the potential range of a primary ZIKV vector, *Aedes aegypti*, including over 30 states [3–5]. Of the 2,487 identified imported ZIKV cases in the US through the end of August, 137 had occurred in Texas. Given historic small, autochthonous outbreaks (ranging from 4 - 25 confirmed cases) of another arbovirus vectored by *Ae. Aegypti*— dengue (DENV) [5–7], Texas was known to be at risk for autochthonous arbovirus transmission, and the recent outbreaks have highlighted the need for increased surveillance and optimized resource allocation in the states and the rest of the vulnerable regions of the Southern United States.

As additional ZIKV waves are possible in summer 2017, public health professionals will continue to face considerable uncertainty in gauging the severity, geographic range of local outbreaks, and appropriate timing of interventions, given the large fraction of undetected ZIKV cases (asymptomatic) and economic tradeoffs of disease prevention and response [8–11]. Depending on the ZIKV symptomatic fraction, reliability and rapidity of diagnostics, importation rate, and transmission rate, the detection of five autochthonous cases in a Texas county, for example, may indicate a small chain of cases from a single importation, a self-limiting outbreak, or a large, hidden epidemic underway (Fig 1). These diverging possibilities have precedents. In French Polynesia, a handful of ZIKV cases were reported by October 2013; two months later an estimated 14,000-29,000 individuals had been infected [8,9]. By contrast, Anguilla had 17 confirmed cases from late 2015 into 2016 without a subsequent epidemic, despite large ZIKV epidemics in surrounding countries [12]. To address the uncertainty, the CDC issued guidelines for state and local agencies; they recommend initiation of public health responses following local reporting of *two* non-familial autochthonous ZIKV cases [13].

**Fig 1.**
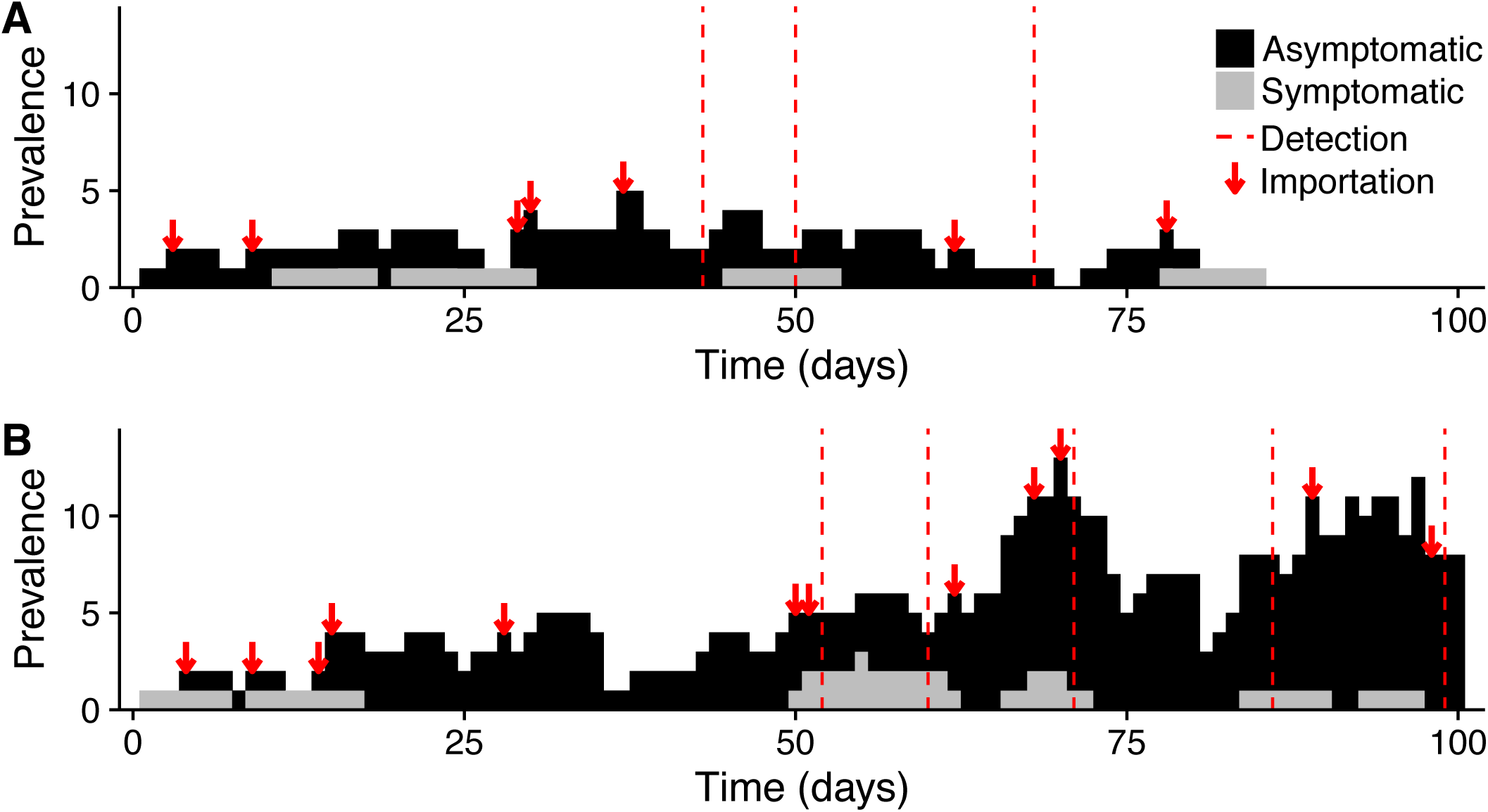
ZIKV emergence scenarios. A ZIKV infection could spark **(A)** a self-limiting outbreak or **(B)** a growing epidemic. Cases are partitioned into symptomatic (grey) and asymptomatic (black). Arrows indicate new ZIKV importations by infected travelers and vertical dashed lines indicate case reporting events. On the 75^th^ day, these divergent scenarios are almost indistinguishable to public health surveillance, as exactly three cases have been detected in both. By the 100^th^ day, the outbreak (A) has died out with 21 total infections while the epidemic (B) continues to grow with already 67 total infections. Each scenario is a single stochastic realization of the model with *R*_*0*_=1.1, reporting rate of 10%, and introduction rate of 0.1 case/day.

Previous risk assessments of ZIKV have provided static *a priori* assessments based on historical incidence and vector suitability, but they do not provide dynamic risk assessments as cases accumulate in a region. Here, we present a framework to support real-time risk assessment, and demonstrate its application in Texas. Our framework accounts for the uncertainty regarding ZIKV epidemiology, including importation rates, reporting rates, local vector populations, and socioeconomic conditions, and can be readily updated as our understanding of ZIKV evolves. To estimate current and future epidemic risk from real-time ZIKV case reports, the model incorporates a previously published method for estimating local ZIKV transmission risk and a new model for estimating local importation risk. Across Texas’ 254 counties, we find that the estimated risk of a locally sustained ZIKV outbreak rises precipitously as autochthonous cases accumulate, and that counties at the southern tip of the Texas-Mexico border and in the Houston Metropolitan Area are at the highest risk for ZIKV transmission. This statewide variation in risk stems primarily from mosquito suitability and socio-environmental constraints on ZIKV transmission rather than heterogeneity in importation rates.

## Methods

Our risk-assessment framework is divided into three sections: (1) county-level epidemiological estimates of ZIKV importation and relative transmission rates, (2) county-specific ZIKV outbreak simulations, and (3) ZIKV risk analysis (Fig S1). To demonstrate this approach, we estimate county-level ZIKV risks throughout the state of Texas for August 2016, given that, by May 2016, Texas experienced dozens of ZIKV importations without subsequent vector-borne transmission.

### Estimating Importation Rates

Our analysis assumes that any ZIKV outbreaks in Texas originate with infected travelers returning from active ZIKV regions. To estimate the ZIKV importation rate for specific counties, we (1) estimated the Texas statewide importation rate (expected number of imported cases per day) for August 2016, (2) estimated the probability *(import risk)* that the next Texas import will arrive in each county, and (3) took the product of the state importation rate and each county importation probability.

1. During the first quarter of 2016, 27 ZIKV travel-associated cases were reported in Texas [5], yielding a baseline first quarter estimate of 0.3 imported cases/day throughout Texas. In 2014 and 2015, arbovirus introductions into Texas increased threefold over this same time period, perhaps driven by seasonal increases in arbovirus activity in endemic regions and the approximately 40% increase from quarter 1 to quarter 3 in international travelers to the US [14]. Taking this as a *baseline* (lower bound) scenario, we projected a corresponding increase in ZIKV importations to 0.9 cases/day (statewide) for the third quarter.
2. To build a predictive model for import risk, we fit a probabilistic model (maximum entropy) [15] of importation risk to 183 DENV, 38 CHIKV, and 31 ZIKV Texas county-level reported importations from 2002 to 2016 and 10 informative socioeconomic, environmental, and travel variables (Supplement §1.1). Given the geographic and biological overlap between ZIKV, DENV and Chikungunya (CHIKV), we used historical DENV and CHIKV importation data to supplement ZIKV importations in the importation risk model, while recognizing that future ZIKV importations may be fueled by large epidemic waves in neighboring regions and summer travel, and thus far exceed recent DENV and CHIKV importations [16]. Currently, DENV, CHIKV, and ZIKV importation patterns differ most noticeably along the Texas-Mexico border. Endemic DENV transmission and sporadic CHIKV outbreaks in Mexico historically have spilled over into neighboring Texas counties. In contrast, ZIKV is not yet as widespread in Mexico as it is in Central and South America, with less than 10 reported ZIKV importations along the border to date (October 2016). We included DENV and CHIKV importation data in the model fitting so as to consider potential future importation pressure from Mexico, as ZIKV continues its increasing trend since March 2016 [17]. To find informative predictors for ZIKV importation risk, we analyzed 72 socio-economic, environmental, and travel variables, and removed near duplicate variables and those that contributed least to model performance, based on out-of-sample cross validation of training and testing sets of data [18,19], reducing the original set of 72 variables to 10 (Tables S3-S4). We validated our importation model by comparing the predicted distribution of cases across the state given a total number of imported cases (September 2016) as a linear predictor of the empirical distribution of cases across counties.

### County Transmission Rates (*R*_*0*_)

The risk of ZIKV emergence following an imported case will depend on the likelihood of mosquito-borne transmission. For emerging diseases like ZIKV, the public health and research communities initially face considerable uncertainty in the drivers and rates of transmission, given the lack of field and experimental studies and epidemiological data, and often derive insights through analogy to similar diseases. For our case study, we estimated county-level ZIKV transmission potential by *Ae. aegypti* using a recently published model [20], that derives some of its key parameters from DENV data. The utility of our framework depends on the validity of such estimates and will increase as our knowledge of ZIKV improves. However, we expect our results to be robust to most sources of uncertainty regarding ZIKV and DENV epidemiology, as they may influence the absolute but not relative county-level risks.

We estimated the ZIKV reproduction number (*R*_*0*_), the average number of secondary infections caused by a single infectious individual in a fully susceptible population, for each Texas county following the method described in Perkins et al [20]. The method calculates *R_0_* using a temperature-dependent formulation of the Ross-Macdonald model, where mosquito mortality rate (μ) and extrinsic incubation period of ZIKV (n) are temperature dependent functions; the human-mosquito transmission probability (b = 0.4), number of days of human infectiousness (c/r = 3.5), and the mosquito biting rate (a = 0.67) are held constant at previously calculated values [20–25]; and the economic-modulated mosquito-human contact scaling factor (m) is a function of county mosquito abundance and GDP data fit to historic ZIKV seroprevalence data [20]. To account for uncertainty in the temperature-dependent functions (the extrinsic incubation period (EIP) and mosquito mortality rate) and in the relationship between economic index and the mosquito-to-human contact rate, Perkins et al. generated functional distributions via 1000 Monte Carlo samples from the underlying parameter distributions. We assume DENV estimates for these temperature-dependent functions, since we lack such data for ZIKV and these Flaviviruses are likely to exhibit similar relationships between temperature and EIP in *Ae. Aegypti* [25]. We used the resulting distributions to estimate *R*_*0*_ for each county, based on county estimates for the average August temperature, mosquito abundance from Kraemer et al [24], and GDP [25]. Our *R*_*0*_ estimates were similar to those reported by Perkins et al. [20] with 95% confidence intervals spanning from 0 to 3.1 (Fig S3). Given this uncertainty, and that our primary aim is to demonstrate the risk assessment framework rather than provide accurate estimates of *R*_*0*_ for Texas, we use these estimates to estimate relative county-level transmission risks (by scaling the county *R*_*0*_ estimates from 0 to 1). In each simulation, we assume that a county’s *R*_*0*_ is the product of its relative risk and a chosen maximum *R*_*0*_. For our case study, we assume a maximum county-level *R*_*0*_ of 1.5 This is consistent with historical arbovirus activity in Texas (which has never sustained a large arbovirus epidemic) and demonstrates the particular utility of the approach in distinguishing outbreaks from epidemics around the epidemic threshold of *R*_*0*_ = 1.

### ZIKV Outbreak Simulation Model

Assuming mosquito-borne transmission as the main driver of epidemic dynamics, to transmit ZIKV, a mosquito must bite an infected human, the mosquito must get infected with the virus, and then the infected mosquito must bite a susceptible human. Rather than explicitly model the full transmission cycle, we aggregated the two-part cycle of ZIKV transmission (mosquito-to-human and human-to-mosquito) into a single exposure period where the individual has been infected by ZIKV, but not yet infectious, and do not explicitly model mosquitos. For the purposes of this study, we need only ensure that the model produces a realistic human-to-human generation time of ZIKV transmission, and the simpler model is more flexible to disease transmission pathways. We fit the generation time of the ZIKV model to early ZIKV Epidemiological estimates, with further fitting details described in Supplemental §2.4.

The resultant model thus follows a Susceptible-Exposed-Infectious-Recovered (SEIR) transmission process stemming from a single ZIKV infection using a Markov branching process model (Fig S4). The temporal evolution of the compartments is governed by daily probabilities of infected individuals transitioning between disease states. New cases arise from importations or autochthonous transmission (Table S5). We treat days as discrete time steps, and the next disease state progression depends solely on the current state and the transition probabilities. We assume that infectious cases cause a Poisson distributed number of secondary cases per day (via human to mosquito to human transmission), but this assumption can be relaxed as more information regarding the distribution of secondary cases becomes available. We also assume infectious individuals are introduced daily according to a Poisson distributed number of cases around the importation rate. Furthermore, *Infectious* cases are categorized into reported and unreported cases according to a reporting rate. We assume that reporting rates approximately correspond to the percentage (∼20%) of symptomatic ZIKV infections [10] and occur at the same rate for imported and locally acquired cases. Additionally, we make the simplifying assumption that reported cases transmit ZIKV at the same rate as unreported cases. We track imported and autochthonous cases separately, and conduct risk analyses based on reported autochthonous cases only, under the assumption that public health officials will have immediate and reliable travel histories for all reported cases [13].

### Simulations

For each county risk scenario, defined by an importation rate, transmission rate, and reporting rate, we ran 10,000 stochastic simulations. Each simulation began with one imported infectious case and terminated either when there were no individuals in either the *Exposed* or *Infectious* classes or the cumulative number of autochthonous infections reached 2,000. Thus the total outbreak time may differ across simulations. We held *R*_*0*_ constant throughout each simulation, as we sought to model early outbreak dynamics over short periods (relative to the seasonality of transmission) following introduction. We classified simulations as either epidemics or self-limiting outbreaks; epidemics were simulations that fulfilled two criteria: reached 2,000 cumulative autochthonous infections and had a maximum daily prevalence (defined as the number of current infectious cases) exceeding 50 autochthonous cases (Fig S6). The second criterion distinguishes simulations resulting in large self-sustaining outbreaks (that achieve substantial peaks) from those that accumulate infections through a series of small, independent clusters (that fail to reach the daily prevalence threshold). The latter occurs occasionally under low *R*_*0*_s and high importation rates scenarios.

To verify that our simulations do not aggregate cases from clear temporally separate clusters, we calculated the distribution of times between sequential cases (Fig S7). In our simulated epidemics, almost all sequentially occurring cases occur within 14 days of each other, consistent with the CDC’s threshold for identifying local transmission events (based on the estimated maximum duration of the ZIKV incubation period) [13].

### Outbreak Analysis

Our stochastic framework allows us to provide multiple forms of real-time county-level risk assessments as reported cases accumulate. For each county, we found the probability that an outbreak will progress into an epidemic, as defined above, as a function of the number of reported autochthonous cases. We call this *epidemic risk*. To solve for epidemic risk in a county following the *x*th reported autochthonous case, we first find all simulations that experience at least *x* reported autochthonous cases, and then calculate the proportion of those that are ultimately classified as epidemics. For example, consider a county in which 1,000 of 10,000 simulated outbreaks reach at least two reported autochthonous cases and only 50 of the 1,000 simulations ultimately fulfill the two epidemic criteria; the probability of detecting two cases in the county would be 10% and the estimated epidemic risk following two reported cases in that county would be 5%. This simple epidemic classification scheme rarely misclassifies a string of small outbreaks as an epidemic, with the probability of such an error increasing with the importation rate. For example, epidemics should not occur when *R*_*0*_=0.9. If the importation rate is high, overlapping series of moderate outbreaks occasionally meet the two epidemic criteria. Under the highest importation rate we considered (0.3 cases/day), only 1% of outbreaks were misclassified.

This method can be applied to evaluate universal triggers (like the recommended two-case trigger) or derive robust triggers based on risk tolerance of public health agencies. For example, if a policymaker would like to initiate interventions as soon as the risk of an epidemic reaches 30%, we would simulate local ZIKV transmission and solve for the number of reported cases at which the probability of an epidemic first exceeds 30%. Generally, the recommended triggers decrease (fewer reported cases) as the policymaker threshold for action decreases, (e.g. 10% versus 30% threshold) and as the local transmission potential increases (*e.g. R_*0*_* = 1.5 versus *R*_*0*_ = 1.2).

## Results

ZIKV importation risk within Texas is predicted by variables reflecting urbanization, mobility patterns, and socioeconomic status (Table S3), and is concentrated in metropolitan counties of Texas (Fig 2A). In comparing the predictions of this model to out-of-sample data from April to September 2016, the model underestimated the statewide total number of importations (81 vs 151), but robustly predicted the relative importation rates between counties (β=0.97, *R*^2^=0.74, p < 0.001). The two highest risk counties‐‐Harris, which includes Houston, and Travis, which includes Austin, have an estimated 27% and 10% chance of receiving the next imported Texas case respectively and contain international airports.

**Fig 2.**
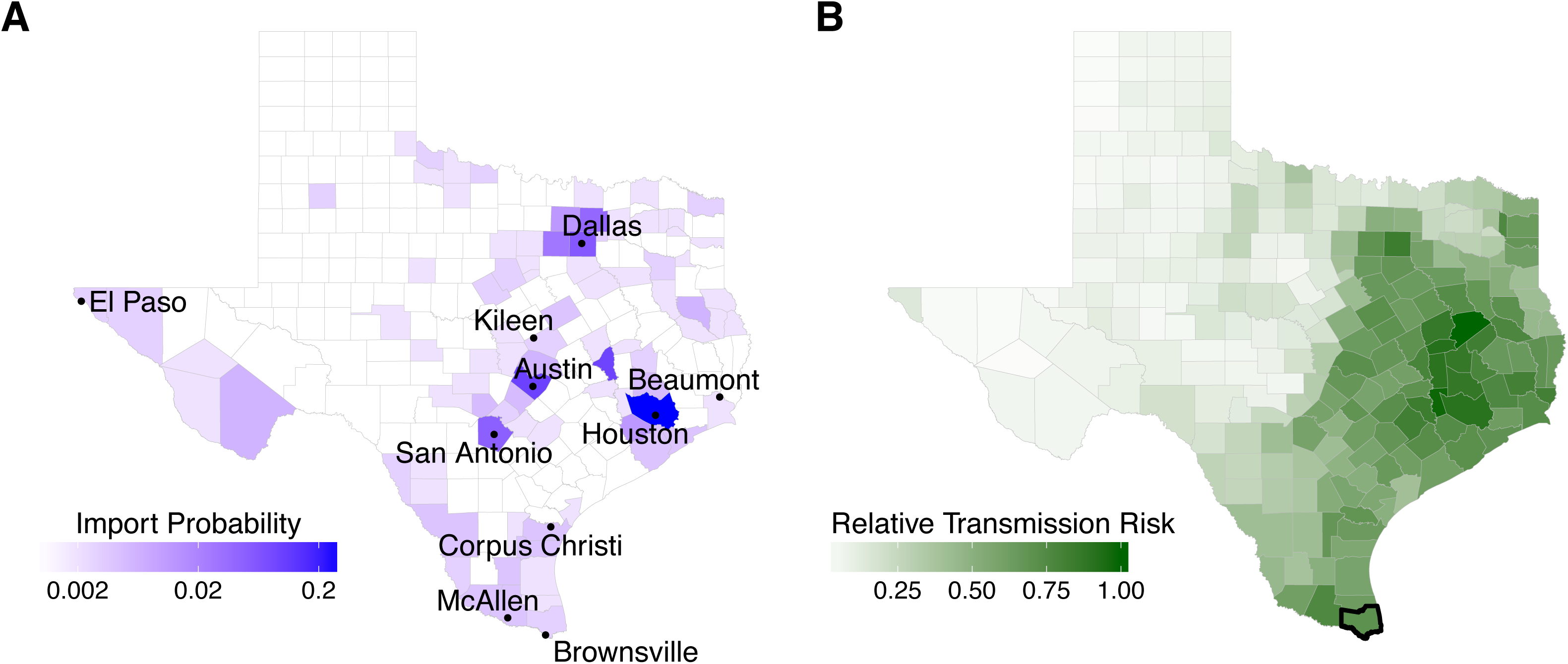
ZIKV importation and transmission risk estimates across Texas for August 2016. (**A**) Color indicates the probability that the next ZIKV import will occur in a given county for each of the 254 Texas counties. Probability is colored on a log scale. The 10 most populous cities in Texas are labeled. Houston’s Harris County has 2.7 times greater chance than Austin’s Travis County of receiving the next imported case. (**B**) Estimated county-level transmission risk for ZIKV (See Fig S7 for seasonal differences). Harris county and Dallas County rank among the top 5 and top 10 for both importation and transmission risk respectively; counties in McAllen and Houston metropolitan area rank among the top 20. Bolded county border indicates counties with recorded local ZIKV transmission.

ZIKV transmission risk is concentrated in southeastern Texas (Fig 2B), partially overlapping with regions of high importation risk (Fig 2A). Our county-level estimates of *R*_*0*_ range widely (from 0.8 to 3.1 for the highest-risk county), reflecting the uncertainty in socioeconomic and environmental drivers of ZIKV (Fig S3). We therefore analyzed the relative rather than absolute transmission risks. For purposes of demonstration, we assumed a plausible maximum county-level *R*_*0*_ of 1.5, which closely followed our median estimates, and scaled the transmission risk for each county accordingly. The following risk analyses can be readily refined as we gain more precise and localized estimates of ZIKA transmission potential.

Wide ranges of outbreaks are possible under a single set of epidemiological conditions (Fig 3A). The relationship between what policymakers can observe (cumulative reported cases) and what they wish to know (current underlying disease prevalence) can be obscured by such uncertainty, and will depend critically on reporting rates (Fig 3B). Under a scenario estimated for Cameron County which experienced the only autochthonous ZIKV transmission in Texas and with a 20% reporting rate, ten linked and reported autochthonous cases correspond to 6 currently circulating cases with a 95% CI of 1-16 from inherent, early-stage outbreak stochasticity. From this wide range of outbreak trajectories, we can characterize time-varying epidemic risk as cases accumulate in a given county. We track the probability of epidemic expansion following each additional reported case in high and low reporting rate scenarios (Fig 3C).

**Fig 3.**
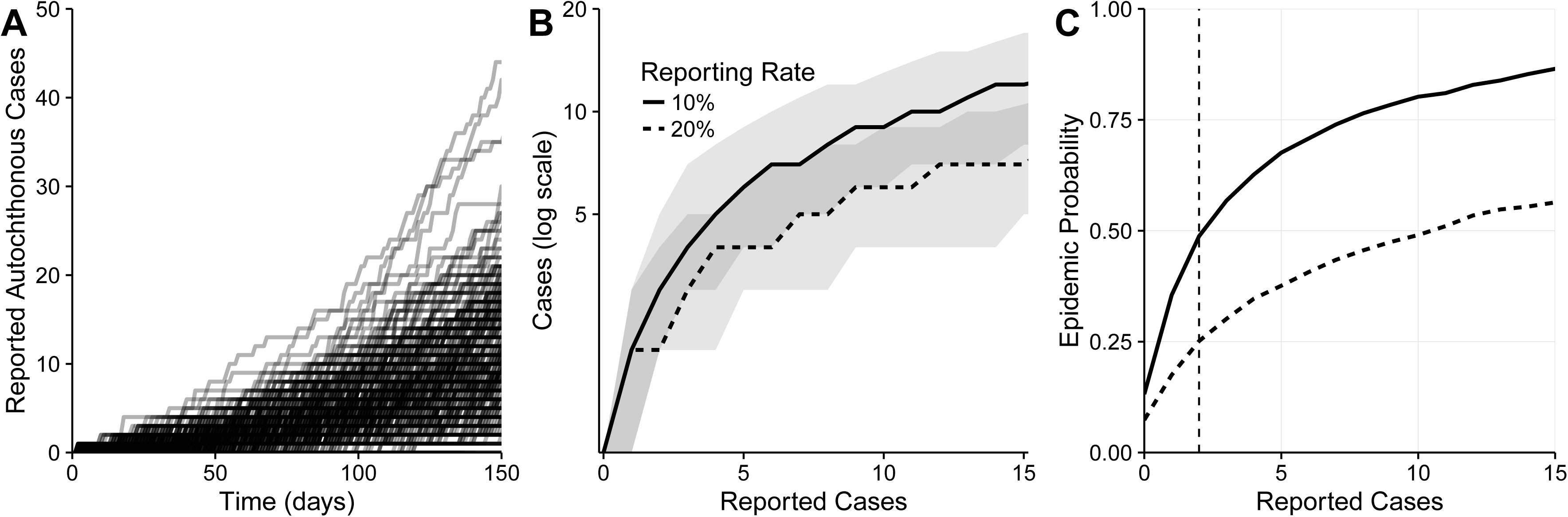
Real-time risk-assessment for ZIKV transmission. All figures are based on transmission and importation risks estimated for Cameron County, Texas. (A) Two thousand simulated outbreaks. (B) Total number of (current) autochthonous cases as a function of the cumulative reported autochthonous cases, under a relatively high (dashed) or low (solid) reporting rate. Ribbons indicate 50% quantiles. (C) The increasing probability of imminent epidemic expansion as reported autochthonous cases accumulate for a low (solid) and high (dashed) reporting rate. Suppose a policy-maker plans to trigger a public health response as soon as a second case is reported (vertical line). Under a 10% reporting rate, this trigger would correspond to a 49% probability of an ensuing epidemic. Under a 20% reporting rate, the probability would be 25%.

These curves can support both real-time risk assessment as cases accumulate and the identification of surveillance triggers indicating when risk exceeds a specified threshold. For example, suppose a policymaker wanted to initiate an intervention upon two reported cases, this would correspond with a 49% probability of an epidemic if 10% of cases are reported, but only 25% if the reporting rate is doubled¨ Alternatively suppose a policy maker wishes to initiate an intervention when the chance of an epidemic exceeds 50%. In the low reporting rate scenario, they should act immediately following the third autochthonous reported case, but could wait until the eleventh case with the high reporting rate.

To evaluate a universal intervention trigger of two reported autochthonous cases, we estimate both the probability of two reported cases in each county and the level of epidemic risk at the moment the trigger event occurs (second case reported). Assuming a baseline importation rate extrapolated from importation levels in March 2016 to August 2016, county *R*_*0*_ scaled from a maximum of 1.5, and a 20% reporting rate, only a minority of counties are likely to experience a trigger event (Fig 4A). While 247 of the 254 counties (97%) have non-zero probabilities of experiencing two reported autochthonous cases, only 86 counties have at least a 10% chance of such an event (assuming they experience at least one importation), with the remaining 168 counties having a median probability of 0.0038 (range 0.0005 to 0.087). Assuming that a second autochthonous case has indeed been reported, we find that the underlying epidemic risk varies widely among the 247 counties, with most counties having near zero epidemic probabilities and a few counties far exceeding a 50% chance of epidemic expansion. For example, two reported autochthonous cases in Harris County, correspond to a 99% chance of ongoing transmission that would proceed to epidemic proportions without intervention, with the rest of the Houston metropolitan also at relatively high risk ranging from 0 (Galveston) to 90% (Waller) (Fig 4B). Given that a universal trigger may signal disparate levels of ZIKV risk, policy makers might seek to adapt their triggers to local conditions. Suppose a policymaker wishes to design triggers that indicate a 50% chance of an emerging epidemic (Fig 4C). Under the baseline importation and reporting rates, an estimated 31 of the 254 counties in Texas are expected to reach a 50% epidemic probability, with triggers ranging from one (Harris County) to 21 (Jefferson County) reported autochthonous cases, with a median of two cases. Counties who detect cases simply due to high importation rates do not have triggers, and the magnitude of a trigger helps quantify a county’s absolute risk for an epidemic as a function of the reported autochthonous cases.

**Fig 4.**
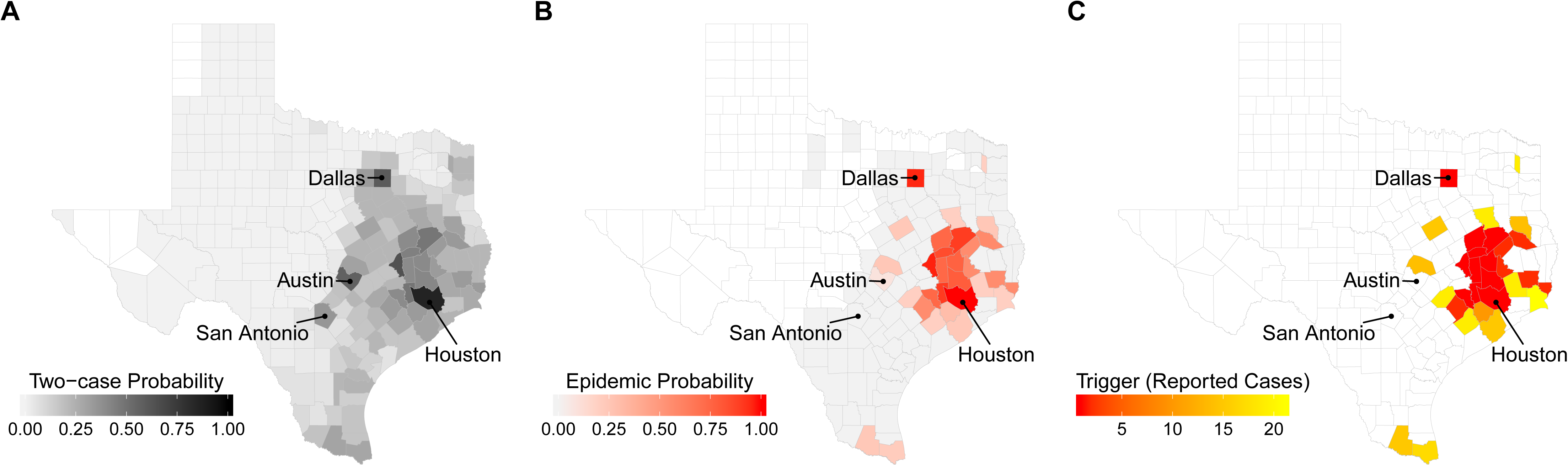
Texas county ZIKV risk assessment. (**A**) Probability of an outbreak with at least two reported autochthonous ZIKV cases. (**B**) The probability of epidemic expansion at the moment the second autochthonous ZIKV case is reported in a county. White counties never reach two reported cases across all 10,000 simulated outbreaks; light gray counties reach two cases, but never experience epidemics. (**C**) Recommended county-level surveillance triggers (number of reported autochthonous cases) indicating that the probability of epidemic expansion has exceeded 50%. White counties indicate that fewer than 1% of the 10,000 simulated outbreaks reached two reported cases. All three maps assume a 20% reporting rate and a baseline importation scenario for August 2016 (81 cases statewide per 90 days) projected from historical arbovirus data. (Figure S5 provides corresponding estimates under a worse case elevated importation scenario).

## Discussion

Our framework provides a data-driven approach to estimating ZIKA emergence risks from potentially sparse and biased surveillance data [26,27]. By mapping observed cases to current and future risks, in the face of considerable uncertainty, the approach can also be used to design public health action plans and evaluate the utility of local versus regional triggers. We demonstrate its application across the 254 ecologically and demographically diverse counties of Texas, one of the two states that has sustained autochthonous ZIKV outbreaks [6,7]. The approach requires local estimates of ZIKV importation and transmission rates. For the Texas analysis, we developed a novel model for estimating county-level ZIKV importation risk and applied published methods to estimate relative county-level transmission risks (Fig 2). We expect that most Texas counties are not at risk for a sustained ZIKV epidemic (Fig 4), and find that many of the highest risk counties lie in the southeastern region surrounding the Houston metropolitan area and the lower Rio Grande valley. However, *R*_*0*_ estimates are uncertain, leaving the possibility that the *R*_*0*_ could be as high as other high risk regions that sustained epidemics [20,28,29]. Our analysis is consistent with historic DENV and CHIKV outbreaks and correctly identifies Cameron county, the only Texas county to have reported local transmission, as a potential ZIKV hot-spot, especially when November estimates are used [30] (Fig S9).

Surveillance triggers‐‐guidelines specifying situations that warrant intervention‐‐are a key component of many public health response plans. Given the urgency and uncertainty surrounding ZIKV, *universal* recommendations can be both pragmatic and judicious. To assist Texas policymakers in interpreting the two-case trigger for intervention guidelines issued by the CDC [13], we used our framework to integrate importation and transmission risks and assess the likelihood and implication of a two-case event for each of Texas’ 254 counties, under a scenario projected from recent ZIKV data to August 2016. Across counties, there is enormous variation in both the chance of a trigger and the magnitude of the public health threat if and when two cases are reported. Given this variation, rather than implement a universal *trigger*, which may correspond to different threats in different locations, one could design local surveillance triggers that correspond to a universal *risk threshold*. Our modeling framework can readily identify triggers (numbers of reported cases) for indicating any specified epidemic event (e.g., prevalence reaching a threshold or imminent epidemic expansion) with any specified risk tolerance (e.g., 10% or 50% chance of that the event has occurred), given local epidemiological conditions. We found close agreement between the recommended two-case trigger and our *epidemic* derived triggers based on a 50% probability of expansion. Of the 30 counties with derived triggers, the median trigger was 2, ranging from one to 21 reported autochthonous cases. These findings apply only to the early, pre-epidemic phase of ZIKV in Texas, when importations occur primarily via travel from affected regions outside the contiguous US.˙

These analyses highlight critical gaps in our understanding of ZIKV biology and epidemiology. The relative transmission risks among Texas counties appear fairly robust to these uncertainties, allowing us to identify high risk regions, including Cameron County in the Lower Rio Grande Valley. Public health agencies might therefore prioritize such counties for surveillance and interventions resources. Given the minimal incursions of DENV and CHIKV into Texas over that past eleven years since the first DENV outbreak in Cameron County, and the high number of importations into putative hotspot counties without autochthonous transmission, we suspect that, if anything, we may be underestimating the socioeconomic and behavioral impediments to ZIKV transmission in the contiguous US. Our analysis also reveals the significant impact of the reporting rate on the timeliness and precision of detection. If only a small fraction of cases are reported, the first few reported cases may correspond to an isolated introduction or a growing epidemic. In contrast, if most cases are reported, policymakers can wait longer for cases to accumulate to trigger interventions and have more confidence in their epidemiological assessments. ZIKV reporting rates are expected to remain low, because an estimated 80% of infections are asymptomatic, and DENV reporting rates have historically matched its asymptomatic proportion [10,31]. Obtaining a realistic estimate of the ZIKV reporting rate is arguably as important as increasing the rate itself, with respect to reliable situational awareness and forecasting. An estimated 8-22% of ZIKV infections were reported during the 2013-2014 outbreak in French Polynesia [29]; however estimates ranging from 1 to 10% have been reported during the ongoing epidemic in Columbia [2,28]. While these provide a baseline estimate for the US, there are many factors that could increase (or decrease) the reporting rate, such as ZIKV awareness among both the public and health-care practitioners, or active surveillance of regions with recent ZIKV cases. Our analysis assumes that all counties have the same case detection probabilities. However, only 40 of the 254 Texas counties maintain active mosquito surveillance and control programs, potentially leading to differences in case detection rates and surveillance efficacy throughout the state [32]. Thus, rapid estimation of the reporting rate using both traditional epidemiological data and new viral sequenced based methods [33] should be a high priority as they become available.

## Conclusions

Our framework can support the development of response plans, by forcing policymakers to be explicit about risk tolerance, that is, the certainty needed before sounding an alarm, and quantifying the consequences of premature or delayed interventions. For example, should ZIKV-related pregnancy advisories be issued when there is only 5% chance of an impending epidemic? 10% chance? 80%? A policymaker has to weigh the costs of false positives‐‐resulting in unnecessary fear and/or intervention‐‐and false negatives‐‐resulting in suboptimal disease control and prevention‐‐complicated by the difficulty inherent in distinguishing a false positive from a successful intervention. The more risk averse the policymaker (with respect to false negatives), the earlier the trigger should be, which can be exacerbated by low reporting rates, high importation rate, and inherent ZIKV transmission potential. In ZIKV prone regions with low reporting rates, even risk tolerant policymakers should act quickly upon seeing initial cases; in lower risk regions, longer waiting periods may be prudent.

## List of abbreviations

ZIKV: Zika virus
DENV: Dengue Virus
CHIKV: Chikungunya Virus
SEIR model: Susceptible-Exposed-Infectious-Recovered epidemiological model
WHO: World Health Organization

## Declarations

### Ethics approval and consent to participate

Not applicable.

### Consent for publication

Not applicable

### Availability of data and materials

The datasets generated and analyzed during the current study are available in the supplemental materials and the R package, rtZIKVrisk, which can be accessed here: https://github.com/sjfox/rtZIKVrisk. There was no individualized patient or medical data used in our study, and all ZIKV case data is publicly available.

### Author Contributions

ND, APG, LAM, LAC, SEB, and SJF developed the conceptual modeling framework and study design. LAC, SJF, and XC collected the data used in the analysis. LAC, KL, and SJF reviewed relevant literature. XC and ND developed and completed the Texas importation risk analysis. LAM, LAC, and SJF interpreted all results. LAC and SJF performed the analyses, created the figures, and wrote the first draft. All authors contributed to the presentation of the results, and the writing and approval of the final manuscript.

## Acknowledgements

We acknowledge the Texas Department of State Health Services (DSHS) for providing historic arbovirus county importation data, and MUG Kraemer and A Perkins for providing Texas mosquito abundance data and technical guidance. We also thank M Johansson and M Meltzer of the Centers for Disease Control and Prevention (CDC) and Dr. John Hellerstedt, commissioner of the Texas DSHS, for critical conversations regarding the modeling framework and its application to Texas and national ZIKV planning efforts. We finally acknowledge the Texas Advanced Computing Center (TACC) at The University of Texas at Austin for providing High performance computing resources that have contributed to the research results reported within this paper. URL:http://www.tacc.utexas.edu.

